# *P. aeruginosa* controls both *C. elegans* attraction and pathogenesis by regulating nitrogen assimilation

**DOI:** 10.1101/2023.11.29.569279

**Authors:** Jacob G. Marogi, Coleen T. Murphy, Cameron Myhrvold, Zemer Gitai

**Affiliations:** Department of Molecular Biology, Princeton University; Lewis Sigler Institute, Princeton University; Department of Chemical and Biological Engineering, Princeton University; Omenn-Darling Bioengineering Institute, Princeton University; Department of Chemistry, Princeton University

## Abstract

Detecting chemical signals is important for identifying food sources and avoiding harmful agents. Like most animals, *C. elegans* use olfaction to chemotax towards their main food source, bacteria. However, little is known about the bacterial compounds governing *C. elegans* attraction to bacteria and the physiological importance of these compounds to bacteria. Here, we address these questions by investigating the function of a small RNA, P11, in the pathogen, *Pseudomonas aeruginosa,* that was previously shown to mediate learned pathogen avoidance. We discovered that this RNA also affects the attraction of untrained *C. elegans* to *P. aeruginosa* and does so by controlling production of ammonia, a volatile odorant produced during nitrogen assimilation. We untangle the complex regulation of *P. aeruginosa* nitrogen assimilation, which is mediated by a partner-switching mechanism involving environmental nitrates, sensor proteins, and P11. In addition to mediating *C. elegans* attraction, nitrogen assimilation is important for bacterial fitness and pathogenesis during *C. elegans* infection by *P. aeruginosa*. These studies define ammonia as a major mediator of trans-kingdom signaling, reveal the physiological importance of nitrogen assimilation for both bacteria and host organisms, and highlight how a bacterial metabolic pathway can either benefit or harm a host in different contexts.

## Introduction

Animals depend on sensory information to promote survival by detecting and responding to a variety of environmental stimuli, such as moving towards nutrient sources or avoiding harmful pathogens. Within their natural environment of soil and rotting vegetation, *Caenorhabditis elegans* are surrounded by diverse bacterial species, which constitute their major food source^1–3^. *C. elegans* respond and chemotax toward volatile compounds using olfactory neurons that control their locomotion^4^. A number of *C. elegans* attractants and repellents have been defined, and some of these attractants and repellants have been shown to be produced by bacteria^4–9^. But the physiological importance of the production of specific bacterially-produced molecules for the chemotaxis of *C. elegans* towards specific bacterial species has remained largely elusive. One example of the complex responses of *C. elegans* to its environment is its multifaceted behaviors in the presence of the bacterium, *Pseudomonas aeruginosa*^8–13^. *Pseudomonas* species represent the most common bacteria found in the *C. elegans* microbiome, suggesting that *C. elegans* encounters these bacteria frequently in nature^3^.

*P. aeruginosa* has a complex regulatory network that controls its pathogenesis such that in some contexts it is nonpathogenic and a beneficial food source for *C. elegans*. But in other contexts, such as higher temperatures or upon association with rigid surfaces, *P. aeruginosa* produces virulence factors that make it pathogenic towards *C. elegans*^14–16^. Many of *P. aeruginosa’s* effects on *C. elegans* are dynamic, as are *C. elegans’* responses to *P. aeruginosa.* Naive *C. elegans* that have never previously encountered *P. aeruginosa* are attracted towards *P. aeruginosa,* and even prefer *P. aeruginosa* over other bacteria like *Escherichia coli* that are constitutively nonpathogenic towards *C. elegans*^10,12,13^. The specific molecule that mediates the attraction of *C. elegans* to *P. aeruginosa* was previously unknown. Here, we identify volatile ammonia, a byproduct of *Pseudomonas aeruginosa* nitrogen assimilation, which mediates *C. elegans* attraction.

Our studies highlight the importance of nitrogen metabolism in *P. aeruginosa*’s interactions with *C. elegans*. Nitrogen is essential for amino acid synthesis and also plays important roles in a variety of other processes^17–19^. Like all animals, *C. elegans* cannot directly utilize inorganic nitrogen sources like ammonium and must therefore obtain processed organic nitrogen from the proteins and amino acids in their bacterial food sources^17,20^. In contrast, bacteria like *Pseudomonas* species can process inorganic environmental ammonium into organic nitrogen^21–24^. Inorganic nitrogen is typically found in two forms: ammonium and nitrate. When ammonium is abundant, it is the preferred nitrogen source and is reduced into glutamine and glutamate, which subsequently function as intracellular organic nitrogen donors^19^. When ammonium availability is limited, most *Pseudomonas* species undergo a process known as nitrogen assimilation where they reduce environmental nitrate into ammonium, which can then be further reduced into glutamine and glutamate^21–24^.

Although many studies have interrogated nitrogen assimilation pathways across diverse bacterial species, its importance in trans-kingdom communication and bacterial host interactions remain largely unknown. Here, we combine *P. aeruginosa* and *C. elegans* genetics to help bridge a gap in trans-kingdom signaling by linking animal chemosensation with bacterial metabolism and physiology. Our data show that volatile ammonia is produced by *P. aeruginosa* nitrogen assimilation, which is regulated by a small RNA (sRNA)-mediated transcriptional termination and antitermination mechanism.

Ammonia produced by *P. aeruginosa* is an attractant for *C. elegans* and causes them to specifically prefer *P. aeruginosa* over *E. coli.* Finally, we found that nitrogen assimilation affects the ability of *P. aeruginosa* to colonize *C. elegans*, which affects pathogenesis and demonstrates the importance of metabolism in trans-kingdom signaling and host-microbe interactions.

## Results

### Disrupting the P11 small RNA perturbs naive attraction of *C. elegans* to PA14

Within their natural habitat, *C. elegans* are surrounded by diverse bacterial species^1–3^. Previous studies have demonstrated that naive, untrained *C. elegans* are attracted to *P. aeruginosa* PA14 (PA14) and prefer it to *E. coli* OP50 (OP50) in a choice assay^10,12,13^. P11 is a small RNA identified in PA14^25^ and later was shown to be necessary for a process in which trained *C. elegans* use transgenerational epigenic inheritance to learn to avoid PA14 for four generations^10,26^. The role of P11 in the attraction of naive *C. elegans* to PA14 has not been previously examined. In the PA14 genome, P11 is found immediately upstream of an operon that mediates the first step of nitrogen assimilation, including the two subunits of nitrite reductase (NirB and NirD), and nitrate reductase (NasC). Given that P11 is upstream of the nitrogen assimilation operon that produces ammonium and that ammonium-related compounds have been previously shown to affect worm chemotaxis^7^, we sought to interrogate P11 function in naive worm chemotaxis.

We examined the role of P11 in mediating the attraction of *C. elegans* to PA14 using an OP50 vs. PA14 bacterial choice assay (Fig. 1a, b). We first confirmed the previous finding that naive *C. elegans* prefer PA14 to OP50. We then performed choice assays between a PA14 strain with a ∼150 base pair deletion surrounding P11 (*ΔP11*) and OP50. We found that *ΔP11* eliminated the preference of PA14 over OP50 (Fig. 1b), suggesting that in the absence of P11, PA14 lack the attractant that mediates preference over OP50.

**Figure 1.**
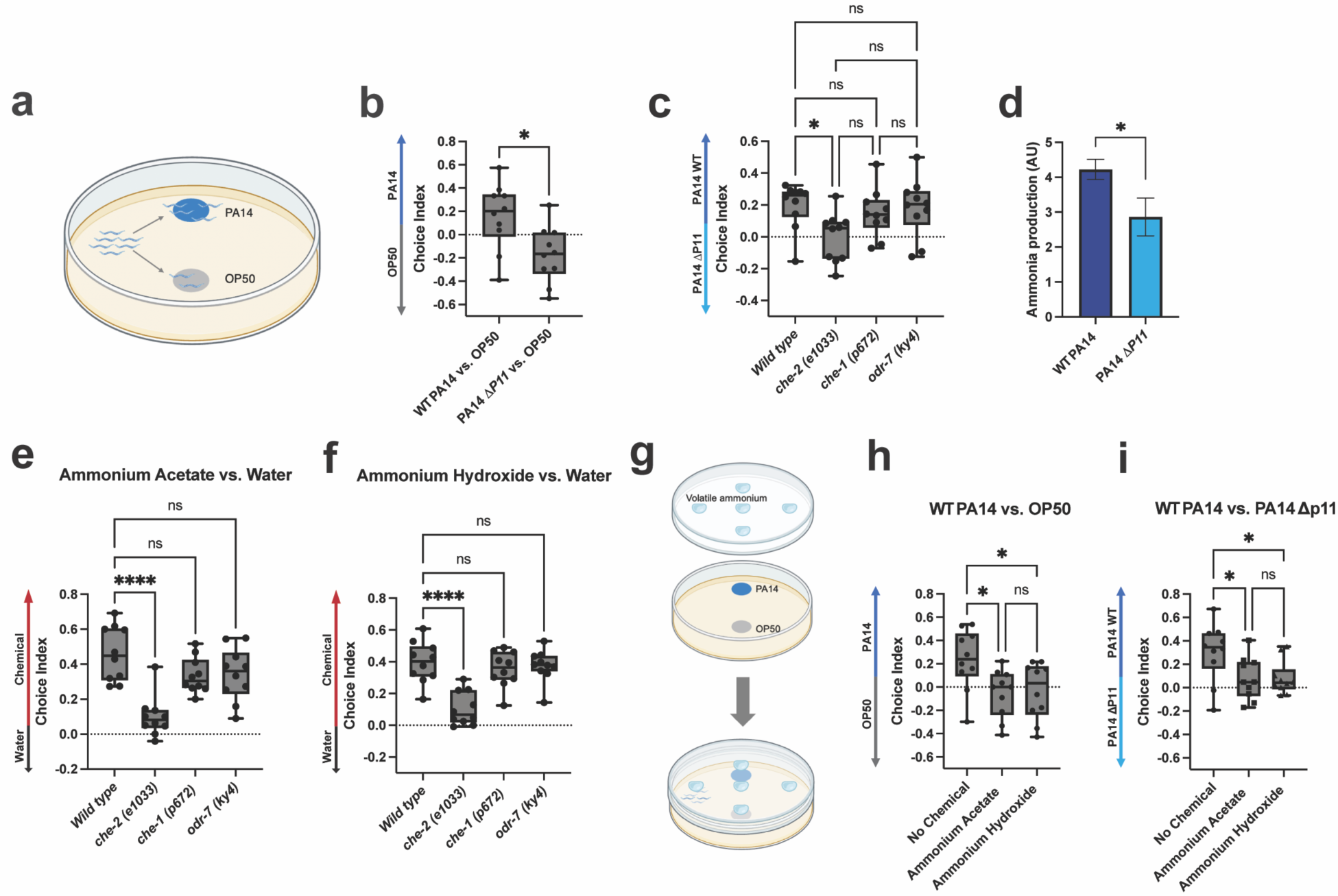
(**a**) Example of worm bacterial choice assay between PA14 and OP50. Worms were hatched and grown on OP50 for 2 days prior to bacterial choice assays. Worms were given 1 hour to chemotax to bacteria and paralyzed at first choice (∼40-100 worms per plate). (**b**) Worm choice assay between PA14 WT*/ΔP11* and OP50. Choice index (CI) = (# on PA14 – OP50)/(Total). (**c**) *che-2(e1033)* worm mutants have defective sensory ciliated neurons and are not naively attracted to PA14 WT. However, *che-1(p672)* and *odr-7(ky4)* worm mutants are ASE and AWA neuron deficient, respectively, and prefer PA14 WT. CI = (# on PA14 WT– PA14 ΔP11)/(Total). (**d**) PA14 WT and ΔP11 ammonia production measured using an ammonia colorimetric assay. Colorimetric fluorescence intensity (ex = 360 nm/ em = 450 nm) normalized to CFU and multiplied by 10^5^. Chemical choice assays between water and ammonium acetate (**e**) and ammonium hydroxide (**f**) with Wild type, *che-2 (e1033)*, *che-1 (p627)*, and *odr-7 (ky4)* worm mutants. CI = (# on chemical - # on water)/(# on chemical + # on water). (**g**) Schematic of bacterial choice assay with saturating concentrations of volatile ammonia. Bacteria spotted on NGM agar and chemical spotted on lid (5 spots of 10 µl chemical, 2.5M each). (**h**) Addition of ammonia (ammonium acetate and ammonium hydroxide) to choice assay between PA14 WT and OP50. (CI) = (# on PA14 – OP50)/(Total) (**i**) Addition of volatile ammonia (ammonium acetate and ammonium hydroxide) to choice assay between PA14 WT and *ΔP11*. CI = (# on PA14 WT– PA14 *ΔP11*)/(Total). Box Plots: center line = median, box range 25^th^ – 75^th^ percentile, minimum/maximum denoted by whiskers, individual choice assay plates are represented by dots. One-Way ANOVA (**b,c,e,f,h,i**) Tukey’s multiple comparison test. Unpaired T-test (**d**). *p≤0.05, ** p≤0.01, *** p≤0.001, **** p≤0.0001, ns = not significant. (Prism 9).

### Worms are attracted to the PA14 nitrogen assimilation metabolite, ammonia

To gain insight into the nature of the P11-dependent signal produced by PA14, we first interrogated the mechanism by which *C. elegans* responds to this signal. Both the AWA and AWC neurons facilitate worm chemotaxis to volatile compounds, whereas the ASE neuron is involved in taste sensation^4,27^. To determine which neuron mediates attraction to PA14, we subjected worms with individual mutants in these neurons to choice assays between wild-type PA14 (WT) and *ΔP11*. Like wild-type *C. elegans,* worms defective in the function of ASE (*che-1(p672)*)^28^ or AWA (*odr-7(ky4)*)^29^ preferred WT PA14 to PA14 *ΔP11*. However, *che-2(e1033)* mutants, which are defective in the function of ciliated neurons including AWA, AWB, AWC, and others^30^, no longer exhibited a preference (Fig. 1c). These results suggest that P11-dependent attraction to PA14 is mediated by a volatile odorant and are consistent with a previous study suggesting that naïve chemotaxis of *C. elegans* to *P. aeruginosa* requires AWC and AWB^13^.

Nitrogen assimilation produces ammonium, which can spontaneously convert to a volatile form, ammonia^31^. Thus, if the deletion surrounding P11 disrupts nitrogen assimilation, it might reduce ammonia production. To determine if the P11 deletion disrupted nitrogen assimilation, we assayed the ability of *ΔP11* to grow in nitrogen-free media supplemented with inorganic nitrates. WT PA14 grew robustly in these conditions, but *ΔP11* mutants did not (Supplementary Fig. 1a). We confirmed that this growth defect was due to an inability to reduce nitrate by demonstrating that adding the products of the nitrogen assimilation reactions (ammonium, Glu, or Gln) rescued PA14 *ΔP11* growth (Supplementary Fig. 1b-e). We also confirmed that the *ΔP11* mutant lacked expression of the downstream nitrogen assimilation genes, providing a molecular explanation for its nitrogen assimilation defect (Supplementary Fig. 1f).

Given that the P11 deletion disrupts nitrogen assimilation, we next sought to directly determine the effect of this mutant on ammonia production. For this purpose, we adapted a colorimetric ammonia assay so that it could be used in the surface-associated conditions employed for *C. elegans* choice assays. We found that *ΔP11* produced ∼2x less ammonia than WT PA14 in the conditions in which we assayed bacterial choice preference (Fig. 1d). We also demonstrated that ammonia is sufficient to promote *C. elegans* attraction by confirming previous findings that *C. elegans* is attracted to ammonia and that this attraction requires the function of the ciliated sensory neurons that detect volatile odorants^7^ (Fig. 1e, f). To determine if ammonia sensing is necessary for *C. elegans’* PA14 preference, we exposed *C. elegans* to saturating levels of exogenous ammonia, which should eliminate any ammonia gradients produced by bacteria in this environment. In the presence of saturating concentrations of ammonia, we found that *C. elegans* no longer preferred WT PA14 to OP50 or *ΔP11* (Fig. 1g-i). Therefore, our data establishes that P11-dependent ammonia production is necessary and sufficient to explain naive *C. elegans’* attraction to PA14.

### A complex regulatory mechanism controls PA14 nitrogen assimilation

The mutant referred to as “*ΔP11”* both above and in prior studies^10^ deletes >150 bp of the genomic region that resides between two operons implicated in nitrogen assimilation (Fig. 2a). We thus examined mutants specific for each of the genes in these operons to dissect their specific functions. As expected, mutating either nitrite reductase, *nirB,* or nitrate reductase, *nasC,* rendered PA14 unable to grow in conditions in which nitrates are the sole nitrogen source (Supplementary Fig. 2a). These mutants also affected *C. elegans* attraction; *C. elegans* preferred WT PA14 to *nirB_Tn_* or *nasC_Tn_.* Loss of *nirB* or *nasC* was also sufficient to completely explain the *ΔP11 C. elegans* attraction defect, as *C. elegans* showed no preference between *ΔP11* and *nirB_Tn_* or *nasC_Tn_* (Fig. 2b).

**Figure 2.**
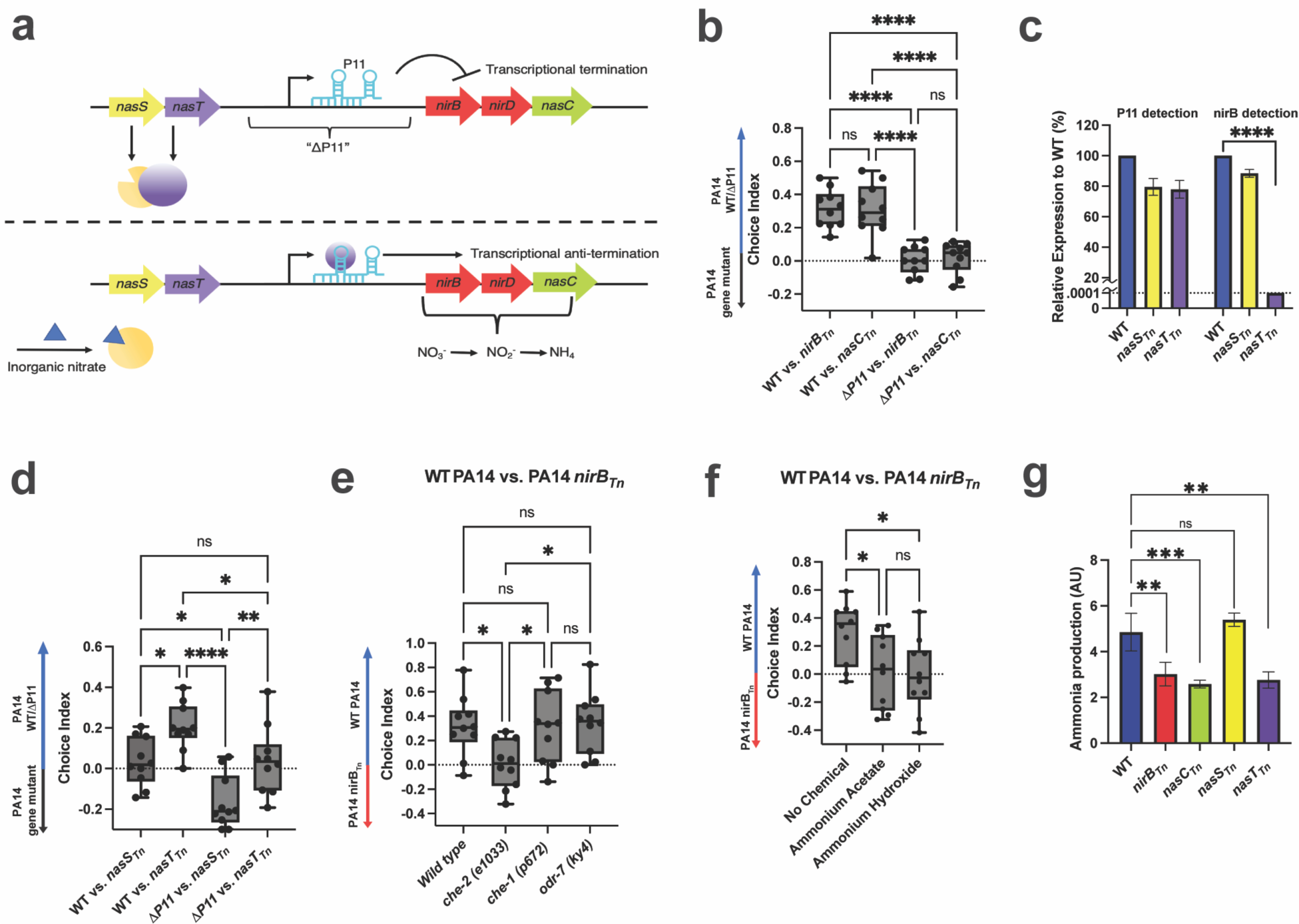
(**a**) Model of PA14 nitrogen assimilation mechanism. When preferred nitrogen sources are limited, the two-component NtrCB system is activated, thus promoting expression at the start of P11. NasS bind to inorganic nitrates and NasT inhibition is relieved. NasT binds to P11, allowing for anti-termination at the leader sequence and expression of the nitrogen assimilation operon. (**b**) Choice assays between PA14 WT*/ΔP11* and PA14 mutants in nitrite reductase (*nirB_Tn_*) and nitrate reductase (*nasC_Tn_*). (**c**) Relative expression of P11 and *nirB* in PA14 *nasS_Tn_/nasT_Tn_* to PA14 WT. Normalized to 5S expression. Dotted line represents limit of detection. (**d**) Choice assays between PA14 WT*/ΔP11* and PA14 mutants in the partner-switching genes (*nasS_Tn_* and *nasT_Tn_*). CI = (# on PA14 WT or *ΔP11* – PA14 gene mutants)/(Total). (**e**) Choice assays between PA14 WT and *nirB_Tn_* with Wild type, *che-2(e1033), che-1(p627)*, and *odr-7(ky4)* worm mutants. CI = (# on PA14 WT – PA14 *nirB_Tn_*)/(Total). (**f**) Addition of ammonia (ammonium acetate and ammonium hydroxide) to choice assay between to PA14 WT and *nirB_Tn_*. CI = (# on PA14 WT – PA14 *nirB_Tn_*)/(Total). (**g**) PA14 WT and gene mutants ammonia production measured using an ammonia colorimetric assay. Colorimetric fluorescence intensity (ex = 360 nm/ em = 450 nm) normalized to CFU and multiplied by 10^5^. Box Plots: center line = median, box range 25^th^ – 75^th^ percentile, minimum/maximum denoted by whiskers, individual choice assay plates are represented by dots. One-Way ANOVA, (**b,d-f**) Tukey’s multiple comparison test, (**c,g**) Dunett’s multiple comparison test. *p≤0.05, ** p≤0.01, *** p≤0.001, **** p≤0.0001, ns = not significant. (Prism 9)

Directly upstream of P11 is another operon containing two regulatory genes, *nasS* and *nasT*. In other *Pseudomonas* strains it has been shown that NasT can bind to either the P11-containing 5’ leader region of the *nirB* gene (P11) or to NasS in a mutually exclusive manner^24^. P11 contains a predicted hairpin that terminates transcription when unbound, and NasT binding to the leader is required for downstream transcription of *nirB, nirD,* and *nasC*. Meanwhile, NasS mutually exclusively binds to either NasT or to nitrate. Thus, in the absence of nitrates, NasS binds to NasT, which leaves P11 unbound and results in termination that prevents *nirBD nasC* expression. When nitrates are present, nitrates bind to NasS, freeing NasT to bind to P11 and promote *nirBD nasC* expression^24^ (Fig. 2a). To determine if a similar partner-switching regulatory mechanism is present in PA14, we assayed PA14 *nasS* and *nasT* mutants for their ability to grow in nitrogen-free media supplemented with nitrates. Whereas *nasS_Tn_* grew as well as WT PA14, *nasT_Tn_* was unable to grow (Supplementary Fig. 2a). We also analyzed the expression of both the P11 and *nirB* RNA’s by qRT-PCR in surface-attached conditions mimicking those used during *C. elegans* choice assays. We found that *nasT_Tn_* dramatically reduced the downstream *nirB* gene to undetectable levels but retained relatively high P11 levels.

While *nasS_Tn_* exhibited similar P11 mutants to those seen in *nasT_Tn_*, the levels of *nirB* were significantly higher in *nasS_Tn_* than in *nasT_Tn_* (Fig. 2c). These results suggest that the NasT-NasS partner-switching mechanism regulates nitrate/nitrite reductase expression in PA14, providing a molecular explanation for their downstream effects on nitrogen assimilation.

Next, we tested the function of *nasS* and *nasT* in *C. elegans* chemotaxis through choice assays against WT PA14 or *ΔP11* and mutants in both genes. Worms demonstrated a preference for WT over *nasT_Tn_* but did not prefer WT to *nasS_Tn_* (Fig. 2d). Conversely, *ΔP11* was less attractive to worms than *nasS_Tn_* but worm attraction to *ΔP11* and *nasT_Tn_* was equal (Fig. 2d). To confirm that mutants in *nirB, nasC,* and *nasT* interfere with *C. elegans* attraction by disrupting the production of a volatile molecule, we assayed *C. elegans* chemotaxis to WT PA14 and mutants in these three genes with the same neuronal mutants as previously described. Worms defective in the function of ASE (*che-1(p672)*)^28^, or AWA (*odr-7(ky4)*)^29^, preferred WT PA14 to *nirB_Tn_, nasC_Tn_,* or *nasT_Tn_*. Meanwhile, worms defective in the function of ciliated sensory neurons (*che-2(e1033)*)^30^, no longer exhibited a preference (Fig. 2e, Supplementary Fig. 2b, c). We further confirmed the identity of the signal as ammonia by introducing exogenous ammonia to choice assays between WT PA14 and *nirB_Tn_, nasC_Tn_,* or *nasT_Tn_*. In the presence of saturating concentrations of ammonia, *C. elegans* were no longer able to distinguish different PA14 strains (Fig. 2f, Supplementary Fig. 2d, e).

Finally, we determined whether our PA14 nitrogen assimilation pathway mutants directly affect ammonia production. Specifically, we assayed ammonia production in surface-attached conditions by PA14 mutants in *nirB, nasC, nasS,* and *nasT.* Mutants that render PA14 unable to assimilate nitrates (*nirB_Tn_, nasC_Tn_, nasT_Tn_*) showed a ∼2x decrease in ammonia production, whereas *nasS_Tn_* produced WT ammonia levels (Fig. 2g). Altogether, these data support our hypothesis that NasS functions as a nitrate sensor and, with NasT, regulates nitrogen assimilation by a partner-switching mechanism. Additionally, all disruptions of the nitrogen assimilation enzymes affect PA14 ammonia production and consequently interfere with *C. elegans* attraction to PA14.

### Two P11 stem-loops inversely regulate nitrogen assimilation expression

RNA secondary structure formation is important for numerous RNA functions and the predicted secondary structure of P11 contains two stem-loops, which often facilitate protein-RNA interactions or function as transcriptional terminators^32–34^. In order to determine the functions of the P11 stem-loops and to better understand the mechanism by which P11 regulates the nitrogen assimilation operon, we made two mutants that each specifically deleted one of the stem-loops (*P11mut1* and *P11mut2*) (Fig. 3a). Unlike the large *ΔP11* deletion described above, these small mutations do not extend into the operon promoter. PA14 *P11mut1* failed to grow in nitrogen-free media supplemented with nitrate (Supplementary Fig. 2f), suggesting that deletion of the first stem-loop leads to a loss of expression of the nitrogen assimilation enzymes. To understand the cause of this nitrogen assimilation phenotypic defect we examined the expression of both the P11 leader region and the downstream *nirB* gene. *P11mut1* retained robust P11 expression, indicating that the operon promoter remained intact. However, *nirB* mRNA expression was so low as to remain undetectable in *P11mut1* (Fig. 3c). These results suggest that *P11mut1* results in transcriptional termination between P11 and *nirB*, which could be explained by loss of NasT binding resulting in a loss of antitermination activity. Conversely, *P11mut2* grew robustly in nitrogen-free media supplemented with nitrate (Supplementary Fig. 2f) and retained relatively high levels of both P11 and *nirB* expression (Fig. 3c). These data suggest that disrupting the second stem-loop leads to constitutive expression of *nirB*. The second P11 stem-loop is followed by a poly-T sequence, a hallmark indicator of a rho-independent terminator^34^. Thus, our results support the hypothesis that within P11, the first stem-loop is the site of antitermination by NasT and the second stem-loop functions as a rho-independent terminator that leads to transcriptional termination when NasT is not bound.

**Figure 3.**
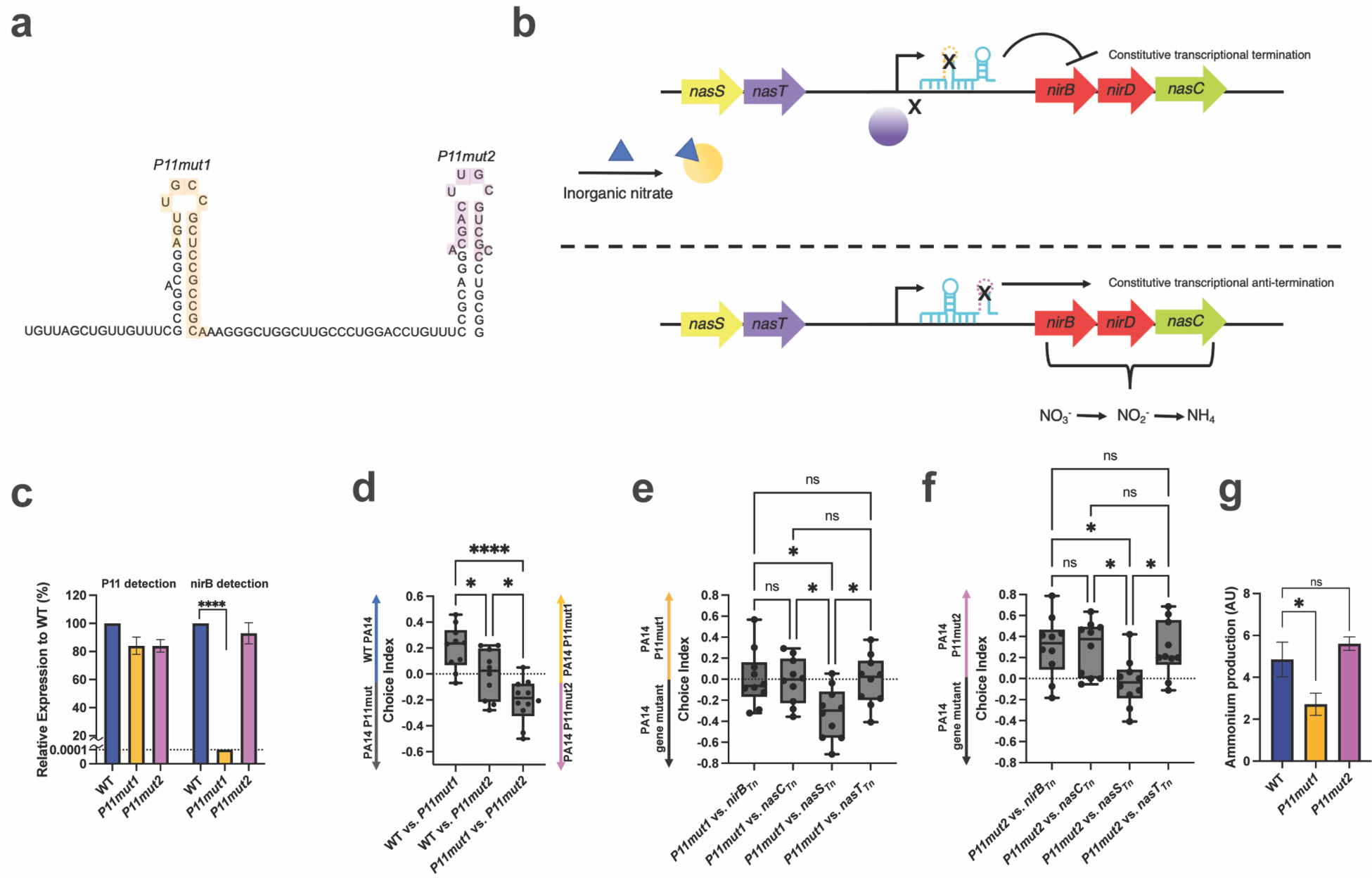
(**a**) Diagram showing putative P11 secondary structure. Yellow: NasT binding site (*P11mut1*) deletion; purple: rho-independent terminator (*P11mut2*) deletion. (**b**) Model showing the effects of *P11mut1* and *P11mut2* on nitrogen assimilation operon expression. P11mut1 prevents NasT binding to P11 and transcriptional anti-termination through the leader sequence (top). *P11mut1* prevents transcriptional termination and therefore, causes constitutive expression of the nitrogen assimilation operon (bottom). (**c**) Relative expression of P11 and *nirB* in PA14 *P11mut1/2* to PA14 WT. Normalized to 5S expression. Dotted line represents limit of detection. (**d**) Choice assays between PA14 WT and *p11mut1/2*. CI = (# on PA14 WT – PA14 P11 mutants)/(Total). Right-hand CI = (# on PA14 *P11mut1* – PA14 *P11mut2*)/(Total) represents last box and whicker plot. (**e**) Choice assays between PA14 P11mut1 and nitrogen assimilation/partner-switching gene mutants. CI = (# on PA14 *P11mut1* – PA14 gene mutant)/(Total) (**f**) Choice assays between PA14 *P11mut2* and nitrogen assimilation/partner-switching gene mutants. CI = (# on PA14 *P11mut2* – PA14 gene mutant)/(Total) (**g**) Ammonia production of P11 mutants. Colorimetric fluorescence intensity normalized to CFU and multiplied by 10^5^. Box Plots: center line = median, box range 25^th^ – 75^th^ percentile, minimum/maximum denoted by whiskers, individual choice assay plates are represented by dots. One-Way ANOVA analysis was performed (**c-g**), Tukey’s multiple comparison test. *p≤0.05, ** p≤0.01, *** p≤0.001, **** p≤0.0001, ns = not significant. (Prism 9).

To examine the phenotypic consequences of stem-loop mutations, we subjected *C. elegans* to choice assays between WT PA14 and the two P11 stem-loop mutants. WT PA14 was preferred over *P11mut1*, whereas no preference was exhibited between WT PA14 and *P11mut2* (Fig. 3d). When given the choice between *P11mut1* and *P11mut2*, *P11mut2* was preferred (Fig. 3d). We also tested worm preference between the P11 mutants and *nirB_Tn_, nasC_Tn_, nasS_Tn_,* and *nasT_Tn_*. Worms did not prefer *P11mut1* over *nirB_Tn_*, *nasC_Tn_*, or *nasT_Tn_* (Fig. 3e). However, *nasS_Tn_* was more attractive than *P11mut1* (Fig. 3e). The opposite effect was observed when comparing mutants in all these genes to *P11mut2*, i.e., worms preferred *P11mut2* over *nirB_Tn_, nasC_Tn_,* and *nasT_Tn_*, but did not prefer *P11mut2* to *nasS_Tn_* (Fig. 3f). Finally, we tested ammonia production in the P11 mutants and found that ammonia levels were reduced in *P11mut1* but were normal in *P11mut2* (Fig. 3g). These results untangle the functions of the two stem-loops of P11 in regulating the expression of the nitrogen assimilation enzymes while providing further support for the central role for P11-dependent ammonia production in mediating the attraction of *C. elegans* to *P. aeruginosa*.

### Nitrogen assimilation is important for PA14 fitness during worm infections

Our findings demonstrate that the nitrogen assimilation pathway affects the attraction of *C. elegans* towards PA14, but a previous study also demonstrated that the large *ΔP11* deletion affects the pathogenesis of PA14 towards *C. elegans*^10^. To determine if the pathogenesis defect of *ΔP11* is due to this mutant’s defect in nitrogen assimilation, we assayed *C. elegans* survival in the presence of a specific mutant in nitrite reductase, *nirB_Tn_*. *C. elegans* infected by *nirB_Tn_* survived significantly longer than worms infected by WT PA14 (Fig. 4a). To confirm that the pathogenesis defect is indeed due to nitrogen assimilation, we also supplemented infected worms with glutamine (a preferred nitrogen source that eliminates the need for nitrogen assimilation). We found that glutamine supplementation fully rescued the pathogenesis defect of *nirB_Tn_*. Thus, nitrogen assimilation is important for optimal pathogenesis of *C. elegans* by PA14.

**Figure 4.**
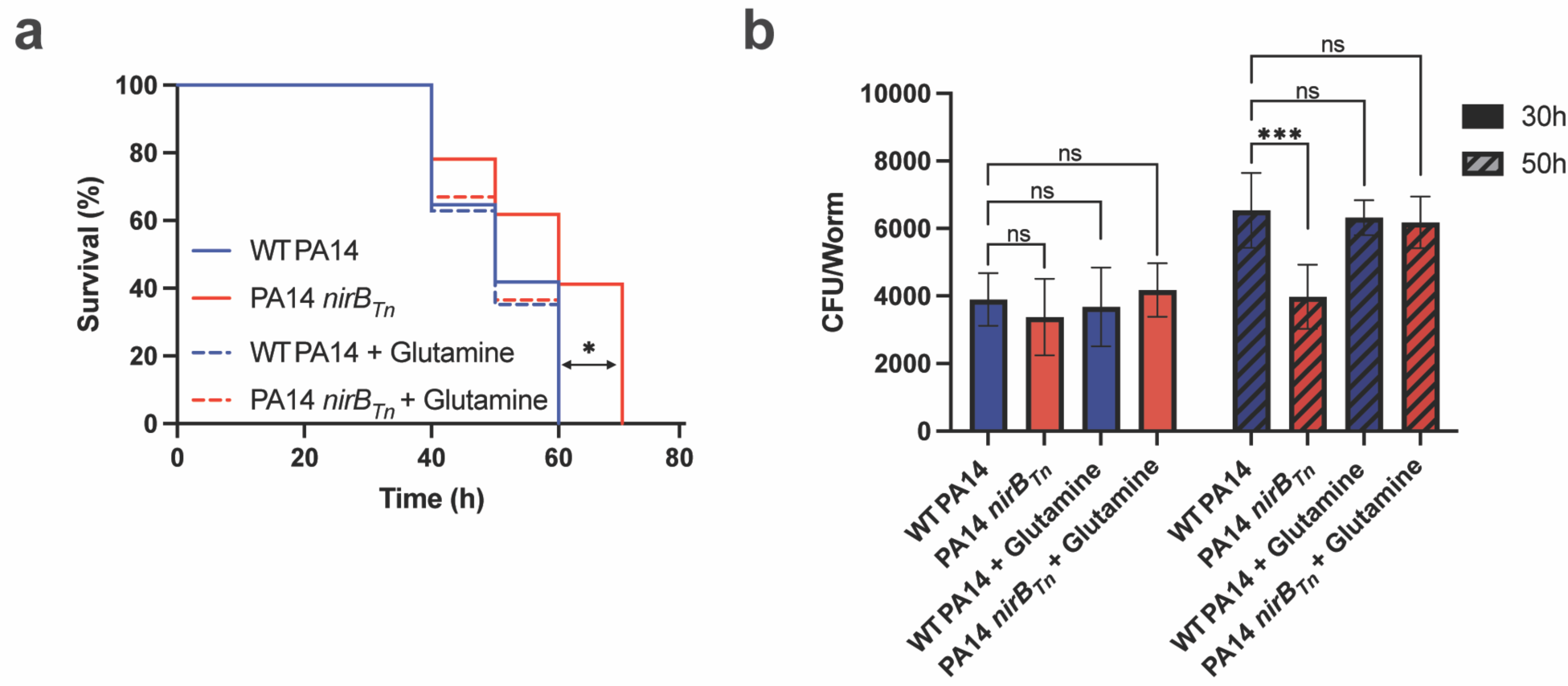
(**a**) Survival of worms infected by PA14 WT and PA14 *nirB_Tn_*. Worms were counted every 10 hours and were declared dead if they were unresponsive to mechanical agitation. Glutamine supplementation (10mM) decreased worm survival (%) infected by PA14 *nirB_Tn_* to survival (%) by PA14 WT infections. P-value calculated using a Mantel-Cox test compared to PA14 WT. (**b**) PA14 CFU’s quantified from worm guts 30h and 50h post infection. At 30h, bacterial load does not differ between strains. At 50h, PA14 *nirB_Tn_* is ∼2x less abundant than PA14 WT and is rescued by glutamine supplementation (10mM). One-Way ANOVA analysis was performed compared to WT. Dunett’s multiple comparison test. *p≤0.05. (Prism 9).

Nitrogen assimilation could influence pathogenesis by affecting the ability of the bacteria to grow within the host (the number of bacteria present), or by affecting the virulence potential of individual bacteria (how harmful each bacterial cell is towards *C. elegans*). To differentiate these possibilities, we assessed the number of bacteria present within *C. elegans* at different timepoints after infection with WT PA14 or *nirB_Tn._* Specifically, we killed all bacteria outside the worms, used mechanical disruption to release the bacteria found inside the worm, and then quantified intrahost bacterial load by quantifying colony forming units (CFU’s) from the contents released from the worms. After 30 hours of infection, we observed no difference in either *C. elegans* survival or intrahost PA14 CFU between WT and *nirB_Tn_* (Fig. 4a, b). However, after 50 hours of infection, we observed that worms survived significantly less in *nirB_Tn_* than in WT, and in these conditions intrahost PA14 CFUs revealed that *nirB_Tn_* was ∼30% less abundant inside of *C. elegans* than WT (Fig. 4b). Supplementing the infections with glutamine, which rescued the survival defect of *nirB_Tn_*, also rescued the intrahost CFU phenotype, restoring the number of *nirB_Tn_* bacteria inside the worms to WT levels (Fig. 4b). Altogether, our data suggest that at later stages of *P. aeruginosa* infection, the environment with *C. elegans* becomes limited for preferred nitrogen sources. Consequently, the ability of these bacteria to assimilate nitrogen becomes important for intrahost growth and colonization in these environments. Mutants that lose the ability to assimilate nitrogen during *C. elegans* infections thus grow more poorly within the host, accumulate to lower levels, and result in less pathogenesis of the host.

## Discussion

Here, we interrogated the functions of nitrogen assimilation in PA14 and how its sRNA-mediated regulation produces volatile ammonia that is attractive to *C. elegans*. *C. elegans* depend on olfactory neurons to sense ammonia from bacteria, which in turn governs their preference towards pathogenic PA14 over less pathogenic PA14 or *E. coli*. Additionally, our data show that PA14 needs nitrogen assimilation for optimal growth and pathogenesis of PA14 during the later stages of *C. elegans* infection.

Our findings demonstrate that the P11 sRNA, which was previously shown to mediate learned avoidance of *C. elegans* by downregulating a specific mRNA in the worm^10^, functions in PA14 as a major regulator of nitrate assimilation. Nitrogen assimilation is an energetically expensive pathway ^19^, which may explain why *P. aeruginosa* PA14 has evolved multiple regulatory components to tightly control its expression. NasS and NasT function in a partner-switching mechanism in response to intracellular inorganic nitrates. By binding to P11, NasT facilitates transcriptional anti-termination of the nitrogen assimilation operon, allowing PA14 to reduce inorganic nitrates into ammonium and subsequently, glutamine and glutamate. Altogether, this multifaceted approach to regulating nitrogen assimilation ensures a proper balance between energy conservation and growth in a specific environment. Interestingly, our results demonstrate that one such environment where nitrogen assimilation is important for bacterial growth is within the *C. elegans* body at later stages of infection. Thus, the pathogenesis defect associated with P11 can be explained by P11’s role in regulating nitrogen assimilation and bacterial growth, rather than by regulating any specific virulence factor that targets *C. elegans*. These data highlight the importance of considering environmental regulation and metabolism in the context of understanding bacterial fitness and pathogenesis during infections. In the future, assaying bacterial load and metabolism within animals may help to untangle the roles of other factors in contributing to either bacterial growth in that environment or to specific virulence pathways.

In addition to shedding new light on the role of P11 in bacteria, our findings provide new insights into how *C. elegans* interact with bacteria in their environment. Some previous studies have examined molecules that are produced by *P. aeruginosa* and sensed by *C. elegans.* These studies identified several compounds that are repulsive to *C. elegans* and one molecule that is attractive, but without being able to eliminate production of these molecules from the bacteria, the actual activity of these molecules remained unclear^8,9,11,35^. Excitingly, we identified the nitrogen assimilation product, ammonia, as the major attractant that causes *C. elegans* to prefer *P. aeruginosa* over *E. coli*. In addition to confirming that ammonia is produced when *P. aeruginosa* attracts *C. elegans,* we demonstrate that ammonia sensing is necessary for *C. elegans’* preference of *P. aeruginosa* over *E. coli.* Specifically, disrupting ammonia gradients by either saturating the environment with excess ammonia or using *P. aeruginosa* mutants defective in ammonia production, inhibits *C. elegans* attraction.

Worms rely on sensory information to find vital resources, such as nitrogen-based compounds, and bacteria provide rich nutrient sources^20,36^. *Pseudomonads* comprise the largest subgroup of bacterial genre within the *C. elegans* natural microbiome^1–3^. Many *Pseudomonads* produce ammonia but are not harmful to *C. elegans,* and many other non-pathogenic soil-based bacteria also possess nitrogen assimilation machinery^3,21–23,37–40^. Therefore, we hypothesize that the benefits of being attracted to nutrient sources signaled by ammonia production may generally outweigh the risk to *C. elegans* of becoming infected by a pathogen. If *C. elegans* does encounter a pathogen like *P. aeruginosa,* pathogenesis is regulated in a complex manner such that the bacteria could be present in a non-pathogenic state^14–16^. And even when *P. aeruginosa* is pathogenic, it often takes several days to kill *C. elegans,* during which time *C. elegans* can still produce offspring. Those offspring will have been taught to move away from *P. aeruginosa* due to the P11-mediated learned avoidance, giving them a chance to survive and find alternative food sources in the next generation^10^. Future work interrogating nitrogen assimilation and ammonia production in different bacteria and how they influence *C. elegans* chemotaxis will shed further light on the generality of these processes.

Our work highlights the complexity of trans-kingdom signaling: the same bacterial pathways that are important for *C. elegans* to acquire essential nutrients are also important for the growth of harmful bacteria within *C. elegans*. The fact that untrained *C. elegans* prefer PA14 suggests that in the short term, the need to find nutrients is prioritized over the risk of encountering a pathogen. However, *C. elegans* have also evolved mechanisms to avoid harmful bacteria in future generations after ingesting them^10,12,26,41^. In this manner, *C. elegans* may use chemotaxis as a simple form of an adaptive immune system that specifically learns to avoid harmful pathogens only once they are encountered. This system of combining ammonia sensing and sRNA-mediated adaptive avoidance, enables *C. elegans* to identify food reservoirs and then avoid pathogens with specificity.

## Material and Methods

Worm strains: Worm strains were gifted by the Murphy lab who received them from the *C. elegans* Genetics Center (CGC) and. N2, *che-2(e1033)* X, *che-1(p672)* I, *odr-7(ky4)* X.

Bacterial strains: OP50 was gifted by the Murphy lab who received it from the CGC. PA14 mutants obtained from a transposon library^42^. The following mutants were used: *nirB::MAR2xT7, nasC::MAR2xT7, nasS::MAR2xT7, nasT::MAR2xT7.* In text mutants are denoted by *Tn* subscript.

General worm maintenance: Worm strains were maintained at 20°C on Nematode Growth Media (NGM) ((3 g/L NaCl, 2.5 g/L Bacto-peptone, 17 g/L Bacto-agar in distilled water, with 1 mL/L cholesterol (5 mg/mL in ethanol), 1 mL/L 1M CaCl2, 1 mL/L 1M MgSO4, and 25 mL/L 1M potassium phosphate buffer (pH 6.0) added to molten agar after autoclaving) plates with *E. coli* OP50 using standard methods.

General bacterial cultivation: OP50 and PA14 were cultured overnight in Luria Broth (10 g/L tryptone, 5 g/L yeast extract, 10 g/L NaCl in distilled water) shaking (250 rpm) at 37°C.

Bacterial cultivation to assay nitrogen assimilation: PA14 strains were grown overnight in LB. 1 mL of overnight culture was washed 2x in minimal salts media (40 mM K2HPO4, 22 mM KH2PO4, 0.5 mM MgSO4, 10 mM FeSO4, pH 7.0) before backdiluting 1:100 in 200 µL minimal salts media supplemented with 20 mM sodium succinate for a carbon source and 10 mM potassium nitrate, ammonium chloride, ammonium sulfate, glutamine, and glutamate for nitrogen sources. OD_600_ readings were performed every 12 minutes for 16 hours in a plate reader.

Bacterial choice assays: Eggs from young hermaphrodites were obtained by bleaching and hatched on NGM plates. Worms were grown at 20°C for 2 days to achieve synchronized L4 populations. Overnight bacterial cultures were diluted to optical density (OD_600_) = 1 and 25 µl were spotted on opposite sides of 60mm NGM plates and incubated for 2 days at 25°C. Plates were incubated at room temperature 1 hour before use and 1 µl of 400mM sodium azide was spotted on each bacteria spot to paralyze worms at first choice. L4’s were washed off of OP50 NGM plates with 1mL M9 buffer and pelleted with gravity for 5 minutes. M9 supernatant was removed and worms were washed again with 1 mL M9 to remove external OP50. After pelleting with gravity for 5 minutes, 5 µL of worms were spotted on choice assay plates (Fig. 1a), and worms were given 1 hour at room temperature to chemotax to bacteria before counted at first choice. ∼40-100 worms were used per plate and 10 plate replicates total were used per condition.

Chemical choice assays: Method as described^7^. Briefly, 10 µL of 5M chemical (Ammonium acetate or ammonium hydroxide) were spotted on 100mm NGM plates opposite of ddH_2_O control immediately before choice assay. L4’s were washed as described above and spotted in the center of the plate between the two choices. Worms were given 1 hour to chemotax to choice and paralyzed with sodium azide (1µL of 400mM sodium azide on each choice). ∼40-100 worms were used per plate and 10 plate replicates total were used per condition.

Bacterial choice assays with excess volatile ammonia: Bacterial choice assay plates and worm preparation were prepared as previously described. Immediately before worms were spotted on choice assay plate, 10 µL spots (5) of 2.5M chemical (ammonium acetate or ammonium hydroxide) were placed on the lid of the plate. Worms were given 1 hour at room temperature to chemotax to bacteria and paralyzed at first choice with sodium azide. ∼40-100 worms were used per plate and 10 plate replicates total were used per condition.

Bacterial RNA isolation: RNA was isolated from PA14 strains grown on NGM plates at 25°C for 2 days. Bacterial cells were isolated from NGM plates by applying 1 mL of M9 buffer and suspending with a cell scraper. Cells were pelleted at 5,000g for 10 min at 4°C. After removing the supernatant, cells were suspended in 1 mL Trizol, vortexed, incubated at 65°C for 10 minutes. Samples were removed and brought to room temperature before adding 200 µL of chloroform, vortexing for 1 minute, and centrifuging at 12,000g for 10 minutes at 4°C. The aqueous phase was inputted for RNA purification using mirVana RNA isolation kit according to manufacturer’s instructions for total RNA. RNA was stored at −80°C.

DNase treatment: 100 µg RNA per sample were treated with DNase according to Invitrogen DNA-free kit manufacturer’s instructions to remove DNA contamination.

qRT-PCR: Two-Step qRT-PCR was performed. First, 500 ng of RNA was used as input for cDNA synthesis. Reverse transcription was performed using a LunaScript RT Supermix Kit according to manufacturer’s instructions. cDNA was diluted 1:10 and 3 µl of diluted cDNA was used as input for qPCR using Luna by NEB master mix for a 20 µL total reaction. 40 cycles were performed for PCR at an annealing temperature of 55°C and extension at 68°C for 20 seconds. For P11 detection primer pair P11.1/2 was used and nirB.1/2 for nirB detection. P11 and nirB expression normalized to 5S expression (primer pair 5S.1/2).

5S.1: GAACCACCTGATCCCTTCCC

5S.2: TAGGAGCTTGACGATGACCT

P11.1: GCGCAAGTTGGCACG

P11.2: CGTTTCCCGACCGAACG

nirB.1: GCGCAAGTTGGCACG

nirB.2: GCCGACCATTACCAGCTTG

Ammonia production assay: 200 µL NGM agar was aliquoted into each well of a flat bottom 96 well plate. Overnight PA14 strains were backdiluted to OD_600_ = 1. 25 µL of culture were spotted in wells and left to dry at room temperature before incubating at 25°C for 2 days. Plates were equilibrated to room temperature for 15 min. Master mix of ammonia colorimetric assay was spotted on bacteria for 15 min, removed by pipetting, and centrifuged at 10,000g for 2 minutes to remove debris before fluorescent intensity measures (all according to manufacturer’s instructions). CFU’s were quantified by resuspending bacteria grown in the same conditions in LB. Fluorescent intensity normalized to CFU to get final relative differences in ammonia production.

P11 mutants strain construction: P11 mutants were constructed by a two-step allelic exchange using plasmid pEXG2. ∼500 bp upstream and downstream of the deletion were amplified from genomic DNA using primer pairs P11mut1-1/2 and P11mut1-3/4 for P11mut1 and P11mut2-1/2 and P11mut2-3/4 for P11mut2. Fragments were fused by overlap extension PCR and inserted by Gibson assembly into pEXG2 (PCR linearized with primers pEXG2-1/2). The pEXG2 plasmid was conjugated and integrated into PA14 genome from donor cells *E. coli* S17. Exconjugates were selected on gent 30µg/mL and irg 100µg/mL and mutants were counterselected on sucrose 15%. Mutants were verified using Seq1/2 primers.

pEXG2-1: AATTAATTTCCACGGGTGCGCATG

pEXG2-2: CTTTACATTTATGCTTCCGGCTCGTA

P11mut1-1: CGCACCCGTGGAAATTAATTGCTATCCCTATGGCGAGATCG

P11mut1-2: TTCAGCATGCTTGCGGCTCGAGTTGGCAGGCGCTTTGTTGT

P11mut1-3: AACTCGAGCCGCAAGCATGCTGAACGTTTCCCGACCGAACGGG

P11mut1-4: CCGGAAGCATAAATGTAAGCTTGAGGCCGTTGGCC

P11mut2-1: CGCACCCGTGGAAATTAATTGCTATCCCTATGGCGAGATCG

P11mut2-2: TTCAGCATGCTTGCGGCTCGAGTTGACGCCTTTGTCCAGGTC

P11mut2-3: AACTCGAGCCGCAAGCATGCTGAAGACGCCTTTTTTGTTTGCGC

P11mut2-4: CCGGAAGCATAAATGTAAGCTTGAGGCCGTTGGCC

Seq1: GCTATCCCTATGGCGAGATCG

Seq2: GCTTGAGGCCGTTGGCC

Survival assay: Wildtype worms were maintained on *E. coli* OP50-coated (NGM) plates prior to experiments. For *P. aeruginosa*-coated plates, overnight cultures were diluted to OD 600 =1, spread onto NGM plates, incubated overnight at 37°C, and equilibrated to 25°C. 10mM glutamine was supplemented into backdiluted cultures before plating on NGM. For virulence assays, synchronized L4 worms were transferred to *P. aeruginosa* plates. Worms were counted at time t=0, 30, 40, 50 and 60 hours to assess viability and were declared dead if they were unresponsive to mechanical agitations. P-values were calculated using a Mantel-Cox test comparing all conditions to WT PA14 (Prism 9).

CFU quantification inside worms: CFU’s of PA14 inside of worms were analyzed as described^43,44^. Briefly, PA14 were allowed to infect *C. elegans* for either 30 or 50 hours. At the relevant timepoints, external bacteria were eliminated by paralyzing worms in 25 mM levamisole to reduce internalization of additional bacteria, placed on NGM plates with 1 mg/mL carbenicillin and 1 mg/mL gentamycin for 15 min, and then moved to fresh NGM plates supplemented with same antibiotics for 30 min to kill external bacteria. Worms were then resuspended in 1 mL M9 and mechanically lysed to release internal bacteria. Lysates were serially diluted in M9 and plated on *Pseudomonas* isolation agar plates. Plates were incubated overnight at 37°C before CFU quantification. Ten worms were used for each CFU quantification with 5 replicates total.

## Acknowledgements

We appreciate training received from Jasmine Ashraf and Renee Seto on work with *C. elegans* and Christopher Guan on qRT-PCR. Funding was provided in part by NIH (1R01AT011963-01 to Z.G. and R21 AI168808-01 to C.M) and CDC (75D30122C15113 to C.M). We thank Dr. Cori Bargmann for helpful comments on the preprint. The opinions, findings, and conclusions are solely the responsibility of the authors and do not necessarily represent the official views of the funding sources.

---

**Supplementary Figure 1.**
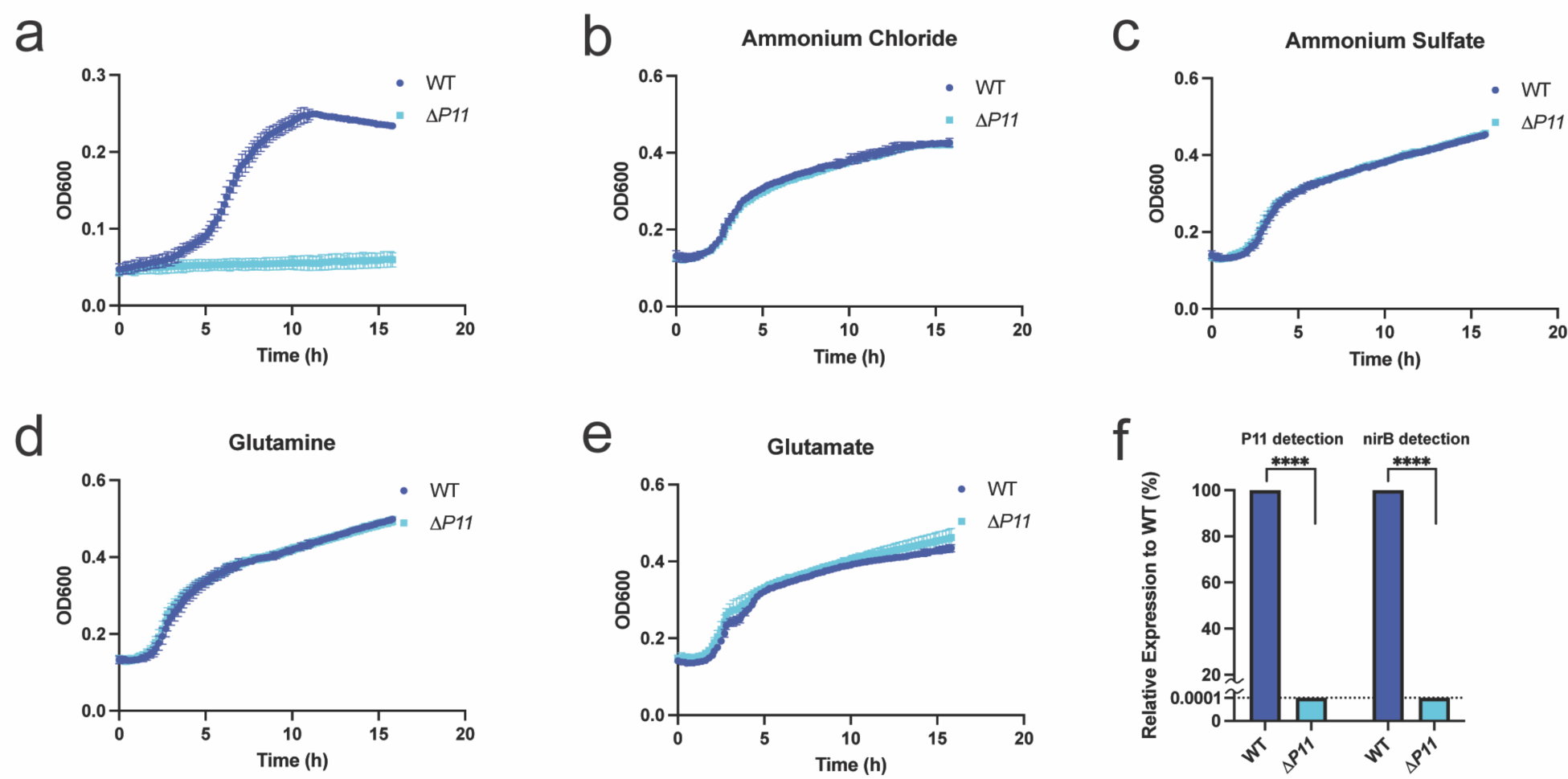
(**a**) PA14 WT and ΔP11 grown at 37°C for 16h in nitrogen free minimal media supplemented with inorganic nitrates. (**b-e**) PA14 WT and *ΔP11* grown 37°C for 16h in nitrogen free minimal media supplemented with preferred nitrogen sources (ammonium chloride, ammonium sulfate, Glu, and Gln, respectively). (**f**) PA14 WT and ΔP11 grown for 48h in surface-attached conditions. Total mRNA collected and *ΔP11* P11 and *nirB* mRNA levels relative to WT calculated. Normalized to 5s expression. Dotted line: limit of detection. (**f**) Unpaired T-test. **** p≤0.0001. (Prism 9).

**Supplementary Figure 2.**
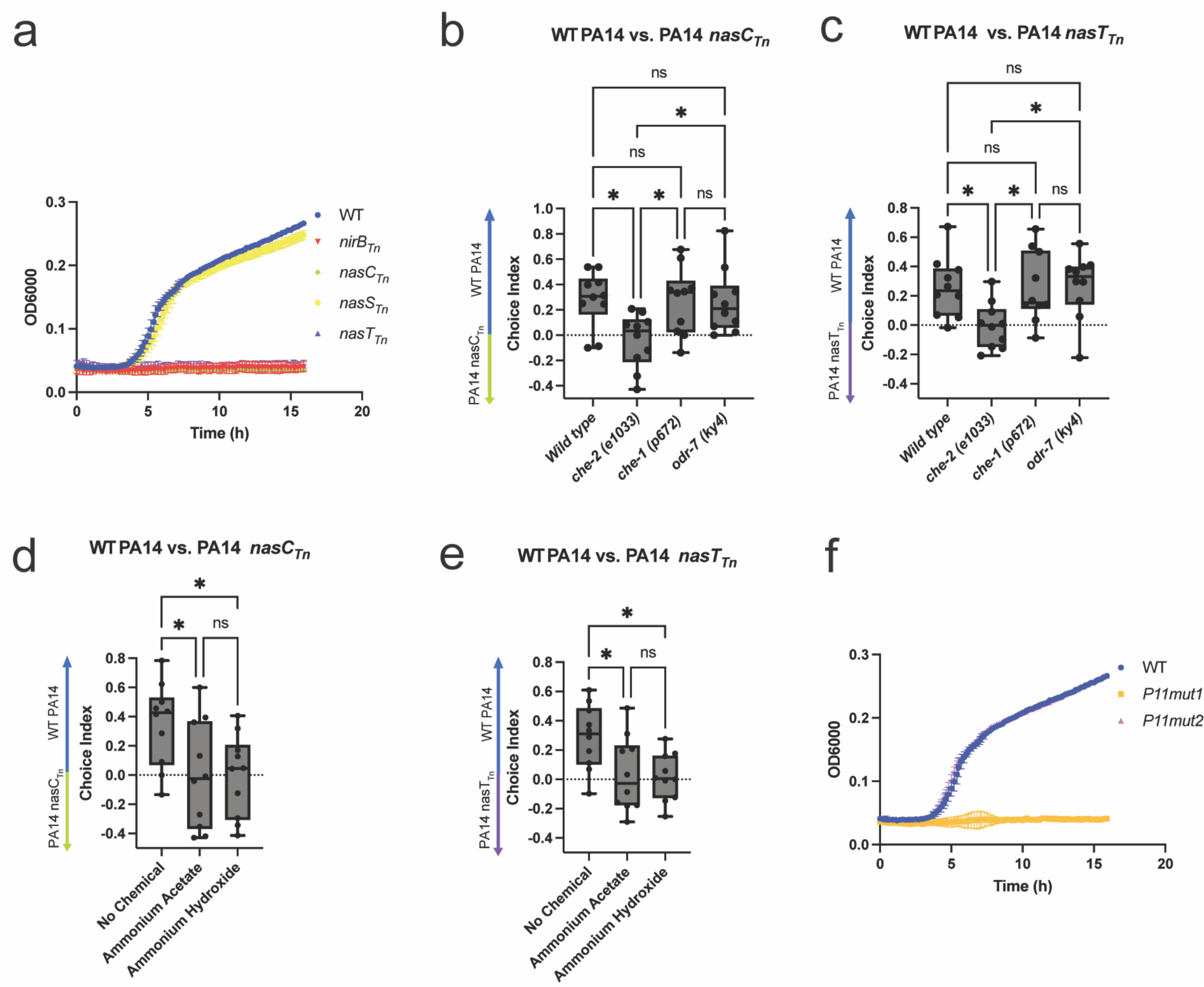
(**a**) PA14 WT, *nirB_Tn_*, *nasC_Tn_*, *nasS_Tn_*, *nasT_Tn_* grown at 37°C for 16h in nitrogen-free minimal media supplemented with inorganic nitrates. (**b**) Wild type, *che-1 (p627)*, and *odr-7 (ky4)* worms prefer PA14 WT over PA14 *nasC_Tn_*, however *che-2 (e1033)* worms showed no preference. CI = (# on PA14 WT – PA14 *nasC_Tn_*)/(Total) (**c**) Wild type, *che-1 (p627)*, and *odr-7 (ky4)* worms prefer PA14 WT over PA14 *nasT_Tn_*, however *che-2 (e1033)* worms showed no preference. CI = (# on PA14 WT – PA14 *nasT_Tn_*)/(Total). (**d,e**) Excess volatile ammonia compounds inhibit worm preference to PA14 WT. CI = (# on PA14 WT – PA14 *nasC_Tn_* or *nasT_Tn_*)/(Total). (**f**) PA14 WT, *P11mut1*, *P11mut2* grown at 37°C for 16h in nitrogen-free minimal media supplemented with inorganic nitrates. One-Way ANOVA analysis was performed (**b-e**), Tukey’s multiple comparison test. *p≤0.05, ** p≤0.01, *** p≤0.001, **** p≤0.0001, ns = not significant. (Prism 9).

